# In-silico and in-vitro evaluation of anti-oxidant and anti-hypertensive activities of *Chenopodium ambrosoides LINN*. ethanol leaf extract

**DOI:** 10.1101/2022.09.22.509042

**Authors:** Ayinde Olaniyi, Oguntoye Oluwatobi, Alabi Oluwabunmi

## Abstract

Oxidative stress and free radicals have been implicated in ethno-pathogenesis of hypertension and cardiovascular disease. *C. ambrosoides* Linn. is a popular plant used in the management of oxidative stress related diseases such as hypertension and obesity in Nigeria and West African countries; however, studies validating the antioxidant and antihypertensive potential of this plant is scanty in literature. Thus, this study was designed to investigate the antioxidant and antihypertensive activities of *C. ambrosoides* ethanolic leaf extract using *in-vitro* (vis-à-vis) DPPH free radical scavenging assay, nitric oxide radical inhibition assay, lipid peroxidation inhibition assay, ferric reducing power assay and angiotensin converting enzyme inhibition assay) and *in-silico* (Molecular docking) techniques and results analyzed using GraphPad prism8 software and Multiple test as criteria for statistical comparison and significance. Gas chromatography mass spectrometry (GCMS) analysis was employed to identify constituent bioactive compounds in the extract. The results of the *in-vitro* anti-oxidants assays show dose dependent inhibition with the highest activity observed at 2.5 mg/ml. The ferric reducing power activity of the extract significantly (P<0.05) shows higher activity than the ascorbic acid standard at all concentration with the highest activity observed at 2.5mg/ml (77.030% against 69.159%).

The extract significantly scavenged DPPH radical than ascorbic acid standard at 2.5mg/ml (81.161% against 75.378%), however at low concentration (1.5mg/ml-0.5mg/ml) the standard shows higher activity than the extract, however dose dependence was maintained. Ascorbic acid standard significantly shows higher activity than the extracts lipid peroxidation inhibition and nitric oxide inhibition activity at all concentration.

The extract exhibited high angiotensin converting enzyme (ACE) inhibited in a dose dependent manner with the highest activity at 2.5mg/ml (95.990%). The ACE inhibitory potential of *C. ambrosoides* extract was corroborated by in-*silico* studies which revealed that 14 out of the 96 identified bioactive compounds through GCMS exhibited higher negative binding affinities than lisinopril (−6.8 Kcal/mol), with the compound 2,4-Diamino-6,8-bis[3,4-dichlorophenyl]-5,6-dihydro-8H-thiapyrano[4’,3’4,5]thieno[2,3d]pyrimidine having the highest binding affinity (−8.0Kcal/mol) In conclusion, it is suggested that the anti-hypertensive activity demonstrated by C. *ambrosoides* might be mediated via its anti-oxidant ability and ACE inhibitory potential.

## 1. Introduction

Checks and balances are the pivot on which maintenance and cellular metabolism and physiology is achieved [1]. Parameters such as temperature, PH, ion concentration, tonicity and water balance, vascular tone and pressure of the biological system are closely monitored as derangement in any of these parameters can lead to partial or total physiological collapse and disarray. Maintenance of normal blood pressure (systolic and diastolic) is of utmost importance, with its deregulation implicated in the development of hypertension and several cardiovascular disorders [2]. Hypertension is a disease characterized by high blood pressure(HBP), and it is a major public health challenge due to its high prevalence around the globe. It is leading cause of morbidity and mortality worldwide with around 7.5 million deaths or 12.8% of the total of all annual deaths worldwide occurs due to high blood pressure and at about 31.1% accounting for 1.39 billion adults worldwide are hypertensive: this figure is projected to proceed to 1.6 billion by 2025, with a persistently elevated blood pressure implicated in the development of stroke, heart failure, ischemic heart disease and renal failure [3].

Apart from genetic factors, the etiology of hypertension is complex. Gender, age, marital status, occupation, BMI, abdominal obesity, and tobacco use have all been pointed as important risk factors in the development of hypertension [4]. The hypothesized pathophysiological mechanisms in the development of hypertension includes but not exclusive to dysregulation of the rennin-angiotensin aldosterone system, endothelial function, mineralocorticoid receptor, vasopressin and aquaporin-2 protein activity and elevated oxidative stress [5].

Synthetic antihypertensive drugs such as Angiotensin converting enzyme–II inhibitors, mineralocorticoids receptor inhibitors and β-channel blockers, and diuretic drugs have been designed over the decade to regulate the elevated blood pressure [3], and in other to make up for the inefficiency of these drugs, the use of combination and chronic usage of anti-hypertensive drugs has been the popular practice, which at that has been reported to elicit different and myriads of undesired side effects such as headache, glucose intolerance, hypotension etc [6]. This informed the need for the development of efficient alternative therapy with less side effect.

Plants are repository of biologically active compounds and have been identified as source of drug development [7]. Furthermore, many medicinal plants are used in folkloric management of cardiovascular diseases and hypertension [8].

*C. ambrosoides* is an annual or perennial plant with a strong aromatic odor that may grow up to 1 meter in height. It is a native herb to tropical Africa and South America, where it is cultivated for use as a leafy vegetable and medicinal herb. The plant has been used to treat influenza, pneumonia, vermicide, gastro-intestinal infections, typhoid, antispasmodic, anti-asthmatic, digestive, vermifuge, emmenagogue, and diabetes in documented articles [9] Moreover, this plant has been reported to contain compounds such as alpha-pinene, aritasone, ascaridole, butyric-acid, d-camphor, essential oils, ferulic-acid, geraniol [10]. However, scientific validation of its pharmacological activity has been relatively understudied. Thus, this study seeks to investigate the antioxidant and antihypertensive activity of C. *ambrosoides* using standard techniques.

## 2. Materials and Methods

### 2.1. Plant material

The fresh sample of *C. ambrosoides* was collected in Ogbomoso town (8° 8’ 31.7940’’ N) by the help of a traditional herb seller. The plant was authenticated by a taxonomist at the department of pure and applied Biology, Ladoke Akintola University of Technology, Ogbomoso, Oyo State, Nigeria

### 2.2. Preparation of ethanol extract

The collected plant was rinsed with clean water to remove sandy particles and thereafter air dried to a constant weight at room temperature for three weeks. The dried plant was pulverized into powder with a kitchen blender (EMEL; EM-242, Shanghai, China). Four hundred (400g) of the pulverized dried plant material was soaked in 1000 ml of ethanol (AnalaR EI/65/457-1) for 72 hours with intermittent shaking. The plant solution (ethanol and plant material) was filtered with a muslin bag and subsequently filtered with filter paper grade 4. The filtrate was thereafter concentrated under pressure at 50°C with a rotary evaporator (Hexa Pharma Chem, India) to obtain a paste-like extract and was refrigerated until further use.

#### 2.2.1. Procedure for serial dilution of plant extract

One thousand milligram (0.1g) of extract was dispensed into 10ml (9 ml H2O and 1 ml of DMSO) of solvent to form the stock solution. Thereafter, serial dilution was prepared for five different concentrations (2.5mg/ml, 2.0mg/ml, 1.5mg/ml, 1.0mg/ml and 0.5mg/ml) from the stock solution in triplicate

### 2.3. GCMS analysis of C. ambrosoides

The GC-MS analysis was performed on Agilent Technologies 789OA coupled with a mass spectrophotometer with triple axis detector (VL5675C) equipped with an auto injector (10 μl syringe). Chromatographic separation was performed on the capillary column (30 m × 250 μm × 0.25 μm) using Helium as the carrier gas at a constant flow rate of 1.5 ml/ min. The sample injection volume was 1 μl in a split mode with a split ratio of 1:50. The column temperature started at 35 °C for 5 min at a rate of 4 ° C/min to 150 °C and was raised to 250 C at the rate of 20 ° C/min with a holding time of 5 min. The total elution time was 47.5 min. The compounds were identified by comparing the spectrum of the separated components with the standard mass spectra from the National Institute of Standards and Technology Library (NIST), Maryland, USA.

### 2.4. Data source for in-silico ACE inhibitory assay

The data for angiotensin converting enzyme (PDB:108A) was obtained from the protein data bank (PDB), (https://www.rcsb.org/structure). The SDS format of the individual ligands from the plants were collected from PubChem database (https://pudchem.ncbi.nlm.nih.gov) and searched on cactus online smiles translator (https://cactus.nci.nih.gov) for ligand download.

Interacting ligands and water molecules were removed from the proteins, thereafter saved in PDB format for docking analysis

### 2.5. Molecular docking and visualization of protein and ligand complex

Docking of the protein with the identified phytochemicals was carried out on Auto Dock Vina 1.5.6. On the Autodock tool. Polar-H-atoms were first added to the proteins followed by Gasteiger charges calculation. The protein file was saved as pdbqt and the grid dimensions were set. Docking calculations and binding affinity (BA) were then performed using Vina folder. Interactions between the ligand and proteins were visualized using discovery studio 2019.

### 2.6. In-vitro anti-oxidant assay

#### 2.6.1. 2,2-diphenyl-1-picrylhydrazyl (DPPH) Assay

The radical scavenging activity of different extract was determined by using DPPH assay according to Chang et al. (2001). The decrease in the absorption of the DPPH solution after the addition of an antioxidant was measured at 517 nm. Ascorbic acid (10mg/ml DMSO) was used as reference.

##### 2.6.1.1. Principle of DPPH

DPPH is a stable (in powder form) free radical with red colour which turns yellow when scavenged. The DPPH assay uses this character to show free radical scavenging activity. The scavenging reaction between (DPPH) and antioxidant (H-A) can be written as,

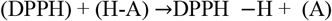

Antioxidant reacts with DPPH and reduces it to DPPH-H and as consequence the absorbance decreases. The degree of discoloration indicates the scavenging potential of the antioxidant compounds or extracts in terms of hydrogen donating ability.

##### 2.6.1.2. Reagent preparation

DPPH (0.1mM) solution was prepared by dissolving 4mg of DPPH in 100 ml of 45% ethanol

##### 2.6.1.3. Procedure of DPPH Assay

Five hundred microliter (500µl) from the stock was measured and poured into a beaker, thereafter, 2.4 mls of the sample was added to 0.6ml of H2O and then 500µl was picked from the mixture into another beaker in triplicate at different concentration (2.5mg/ml, 2.0 mg/ml, 1.5 mg/ml, 1.0 mg/ml and 0.5 mg/ml). Five hundred (500µl) of DPPH solution was added and the mixture was placed in a dark cupboard to incubate for 30 minutes. Thereafter, the absorbance of each mixture was measured at 517 nm. Three (3ml) of DPPH solution was added to the mixture without the extract and was taken as control. The % control scavenging activity of the plant extract was calculated using the following formula;

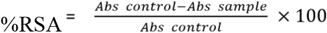

Where RSA is the Radical Scavenging Activity: Abs control is the absorbance of DPPH radical. *Abs* sample is the absorbance of DPPH radical + plant extract.

### 2.7. Reducing power assay

The reducing capacity of extracts was measured using the potassium ferricyanide reduction method. Various concentrations of extract and standard (25-500ug mL-1) were added to 2.5mL of (0.2M) sodium phosphate buffer (pH 6.6) and 2.5 mls of potassium ferricyanide [k_3_Fe_3_(CN)_6_]. One percent solution and vortexed. After incubation at 50ºC for 20min, 2.5ml of Trichloro acetic Acid (TCA) (10%, w/v) was added to all the tubes and centrifuged (Remi, India) at 3000 x g for 10min. Afterwards, upper layer of the solution (5 mls) was to this, 1 mls of FeCI_3_ (1%) was added to each test tube and incubated at 35ºC for 10min. The absorbance of the organic upper layer was measured at 532nm.

### 2.8. Nitric oxide radical (NO^-^) scavenging activity

NO generated from sodium nitroprusside (SNP) was measured according to the method of Marcocciet a. (1994). Briefly, the reaction mixture (5.0 ml) containing SNP (5 mM) in phosphate-buffered saline (pH 7.3), with or without the plant extract at different concentrations, was incubated at 25ºCfor 180 minutes in the presence of a visible polychromatic light source (25W tungsten lamp). The NO radical thus generated interacted with oxygen to produce the nitrite ion (NO.) which was assayed at 30 minutes’ intervals by mixing 1.0ml of incubation mixture with an equal amount of Griss reagent (1% sulphanilamide in 5% phosphoric acid and 0.1% naphthalene-diamine dihydrochloride). The absorbance of the chromophore (purple azo dye) formed during diazotization of nitrite ions with sulphanilamide and the subsequent coupling with naphthalene-diamine dihydrochloride was measured at 546 nM. The nitrite generated in the presence or absence of the plant was estimated using a standard curve based on sodium nitrite solution of known concentrations. Each experiment was carried out in triplicates

### 2.9. Angiotensin converting enzyme (ACE) inhibition assay

Method described by Jimsheena and Gowda, 2003 was used to perform ACE inhibitory assay. The assay mixture contained 125µl of 0.05M sodium borate buffer (pH 8.2) containing 0.3M Sodium Chloride (NaCl), 50µl of 5mM hippuryl-histidyl-leucine (HHL), 25µl of ACE enzyme, all pre-incubated with different samples concentration of the plant inhibitor at 37°C for 30 mins. After stopping the reaction, 0.4ml of pyridine was added followed by 0.2ml of benzene sulphonyl chloride (BSC) and the solution was mixed on a vortex mixer and cooled on ice. The solution was transferred to the 96-well plate. The absorbance was measured at 410nM. The experiments were conducted in triplicate and the following equation was used to calculate the %ACE inhibition.

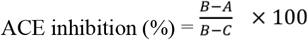

Where B is the absorbance with ACE and HHL without the ACE inhibitor component; A is the absorbance with ACE, HHL and C is the absorbance with HHL without ACE and ACE inhibitor component.

### 2.10. Statistical Analysis

Data obtained were expressed as mean ± SEM and subjected to multiple t-test, significant difference was taken at p < 0.05 using graphPad prism version 8.0. standard curve based on sodium nitrite solution of known concentrations. Each experiment was carried out in triplicates

## 3. Results

### 3.1. In-vitro antioxidant activities of C. ambrosoides extract

#### 3.1.1. DPPH inhibition assay

The DPPH free radical scavenging ability of *C. ambrosoides* with respect to ascorbic acid standard - at five differing concentrations (2.5mg/ml, 2.0mg/ml, 1.5mg/ml, 1.0mg/ml, 0.5mg/ml) is represented in and Figure 1

**Figure 1.**
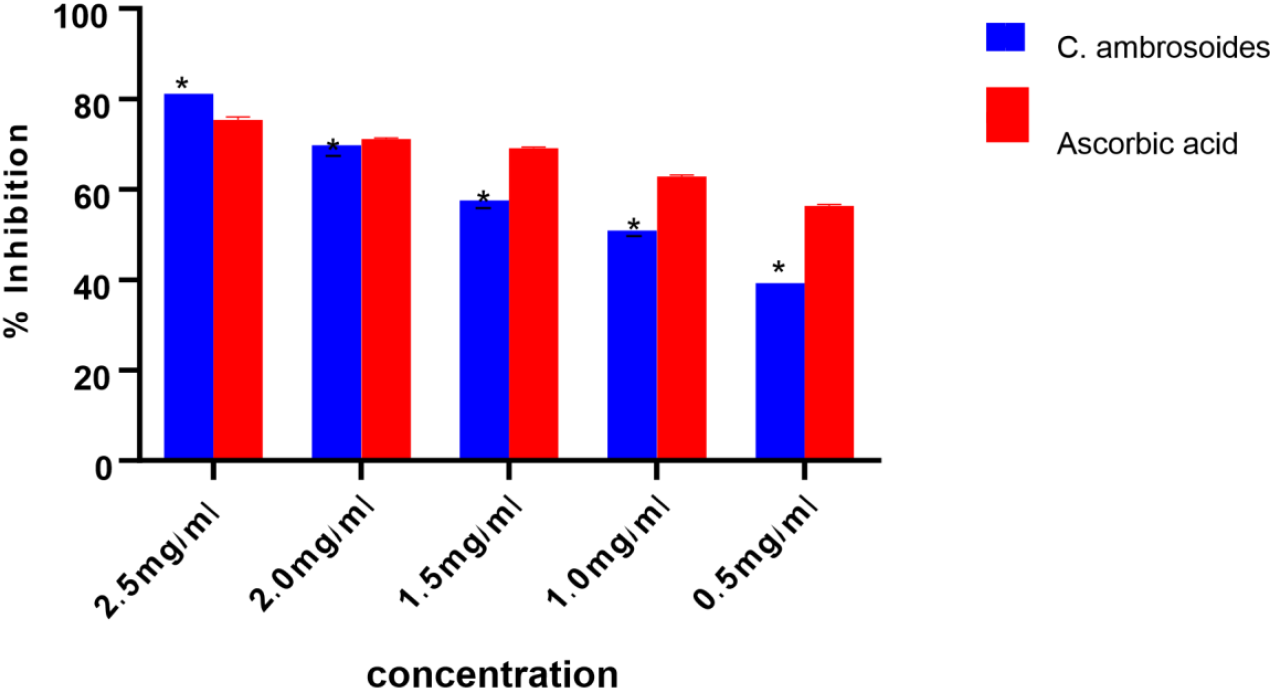
DPPH inhibitory activity of *C. ambrosoides*. Bars with * are significantly different (p<0.05). bars are mean ± standard error mean.

*C. ambrosoides* shows a dose dependent DPPH free radical scavenging activity most potent at 2.5mg/ml (81.161± 0.334), a value significantly higher than observed in standard (75.378 ± 0.666), thus shows higher activity than in the standard at highest concentration but ranked below standard in activity at lower concentrations (1.5mg/ml-0.5mg/ml).

#### 3.1.2. Ferric reducing power assay

The ferric metal (Fe 2+) ions reducing power ability of *C. ambrosoides* with respect to ascorbic acid standard-both at five differing concentrations (2.5mg/ml, 2.0mg/ml, 1.5mg/ml, 1.0mg/ml, 0.5mg/ml) is depicted in Figure 2

**Figure 2.**
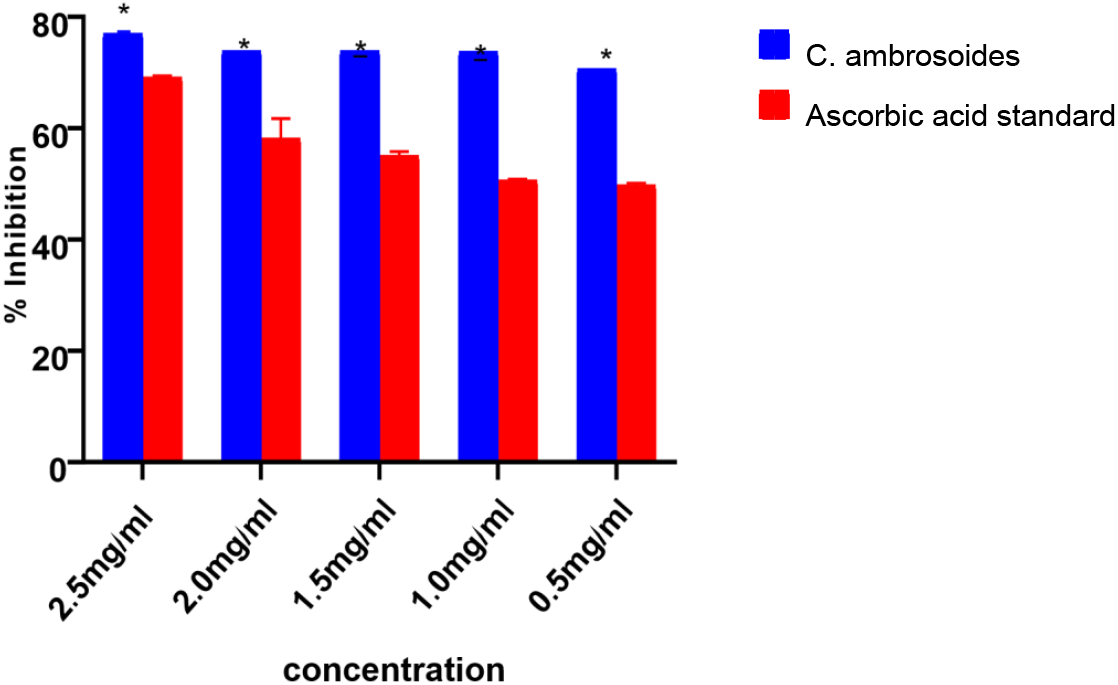
Reducing power activity of *C. ambrosoides*. Bars with * are significantly different (p<0.05). bars are mean ± standard error of mean.

*C. ambrosoides* dose dependently reduce ferric ion, having its highest % metal reducing activity at 2.5mg/ml (77.035± 0.227), a value significantly higher than observed in standard (69.159 ± 0.133). The extract shows higher activity than the standard at all concentrations, although the extract activity comparison across concentration is not significant

#### 3.1.3. Nitric oxide inhibitory activity of C. ambrosoides

The nitric oxide inhibitory activity of *C. ambrosoides* with respect to ascorbic acid standard - at five differing concentrations is shown in Figure 3

**Figure 3.**
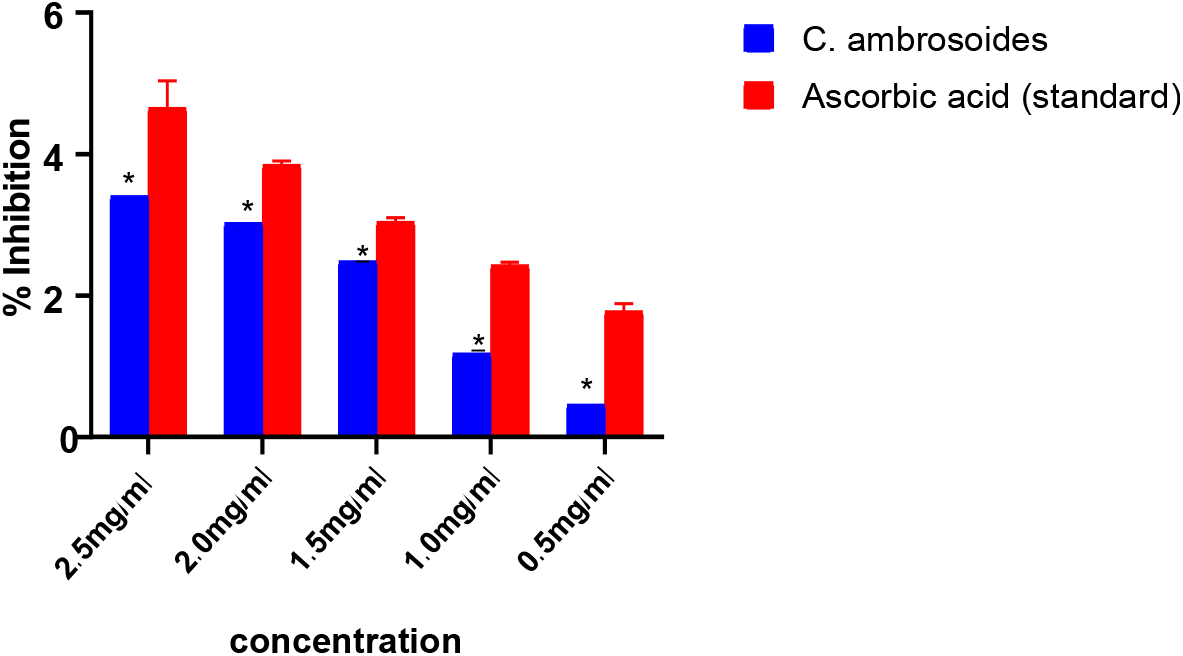
Nitric oxide inhibitory activity of *C. ambrosoides*. Bars with * are significantly different (p<0.05). bars are mean ± standard error of mean.

*C. ambrosoides* shows a dose dependent NO inhibitory activity highest at 2.5mg/ml (34.089± 0.377), a value significantly lower than observed in standard (46.526 ± 2.189). The standard significantly shows higher activity than the extract at all concentration.

#### 3.1.4. Lipid peroxidation inhibitory activity of C. ambrosoides extract

Figure 4 is the representation comparing lipid peroxidation inhibition ability of *C. ambrosoides* at five differing concentrations (2.5mg/ml-0.5mg/ml). *C. ambrosoides* has its highest % lipid peroxidation inhibition activity at 2.5mg/ml (28.279 ± 1.084), a value significantly lower than observed in standard (57.322 ± 0.778) at same concentration. A dose dependence is discovered in both extract and standard, although the standard shows higher activity than the extract at all concentration.

**Figure 4.**
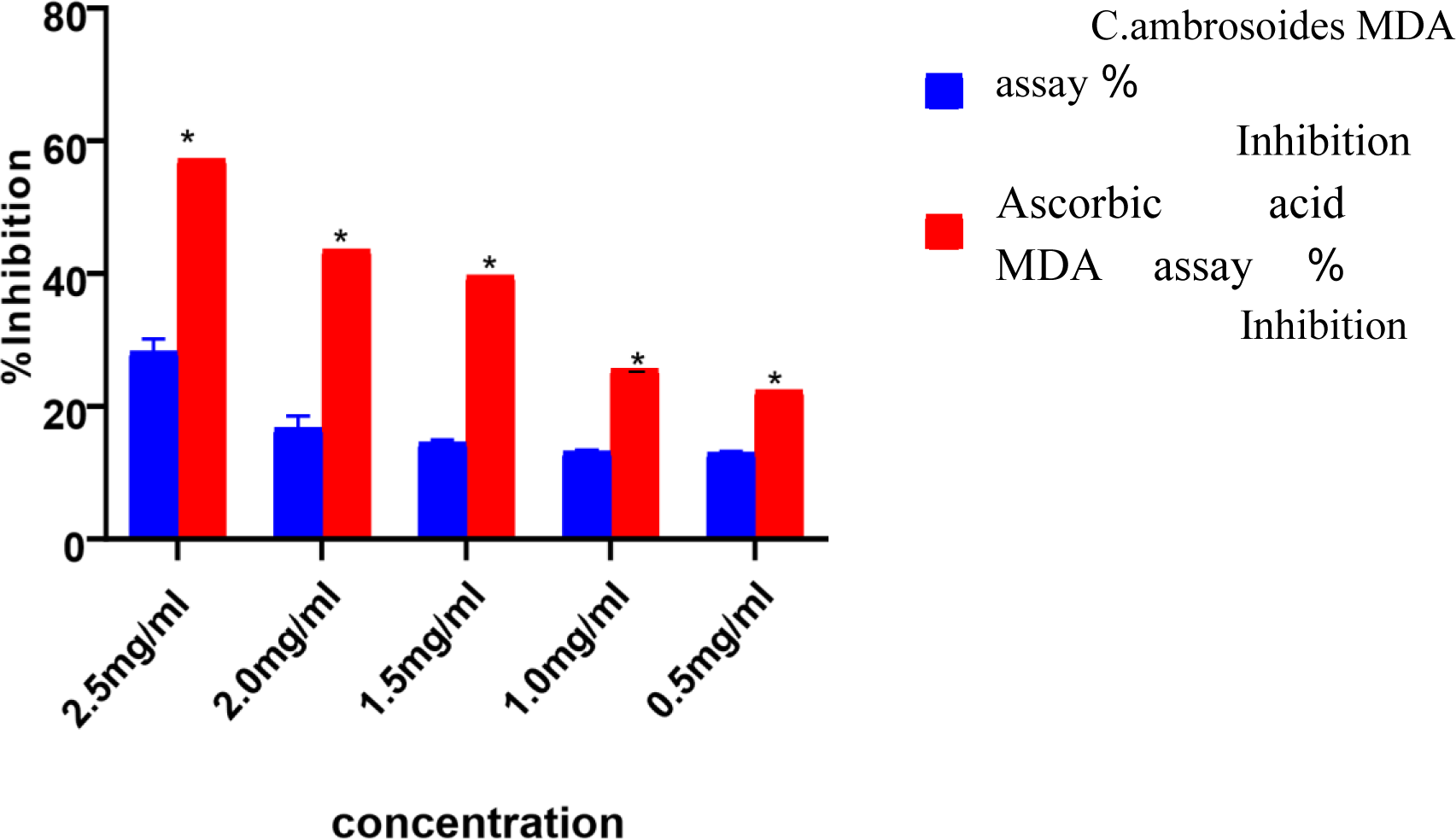
Percentage (%) lipid peroxidation inhibition of C. ambrosoides. Bars with * are significantly different (p<0.05). bars are mean ± standard error of mean.

### 3.2. In- vitro angiotensin converting enzyme inhibition assay

Represented in figure 5 is the percentage angiotensin converting enzyme (ACE) inhibiting activity of *C. ambrosoides* at five differing concentrations (2.5mg/ml, 2.0mg/ml, 1.5mg/ml, 1.0mg/ml, 0.5mg/ml).

**Figure 5.**
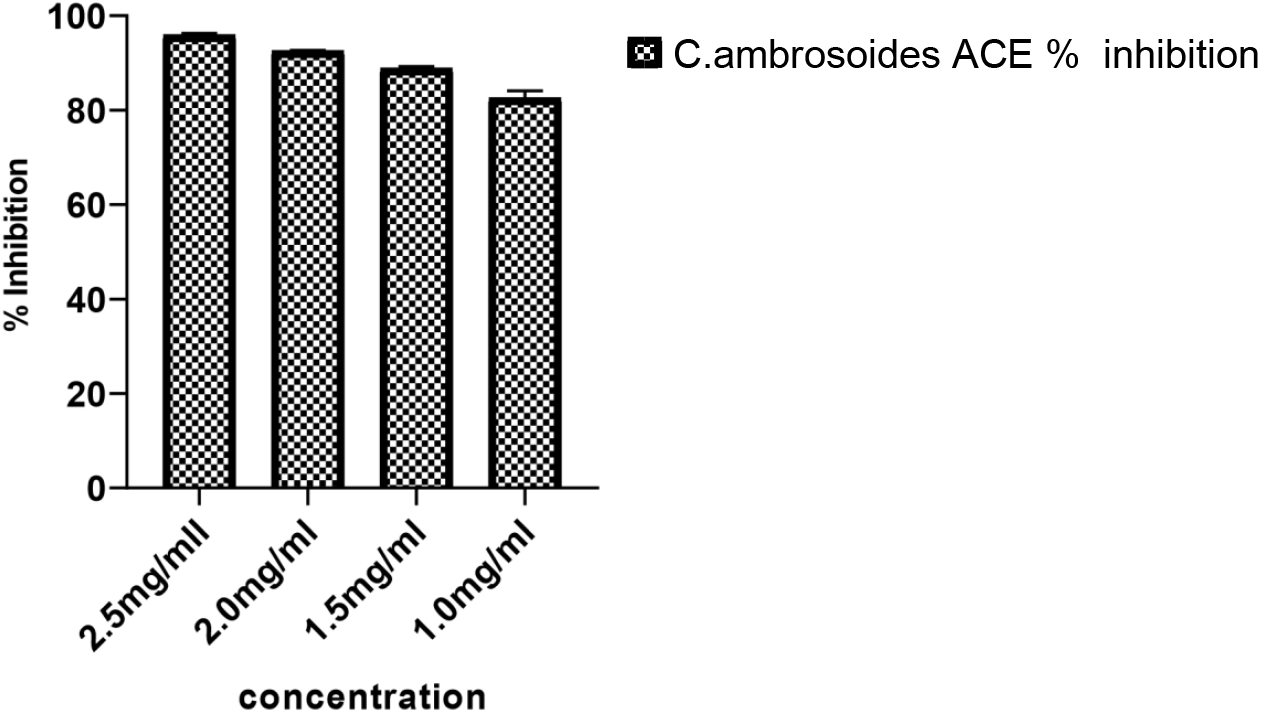
Percentage% ACE inhibitory activity of C. ambrosoides. Bars are mean ± standard error of mean.

**Figure 6.**
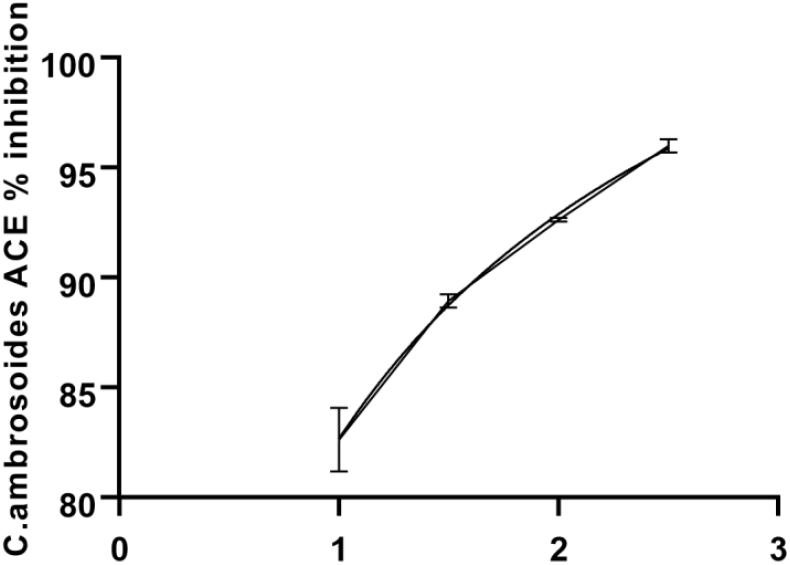
IC50 plot of ACE inhibitory activity of C. *ambrosoides*. IC50 =1.105mg/ml

**Figure 7.**
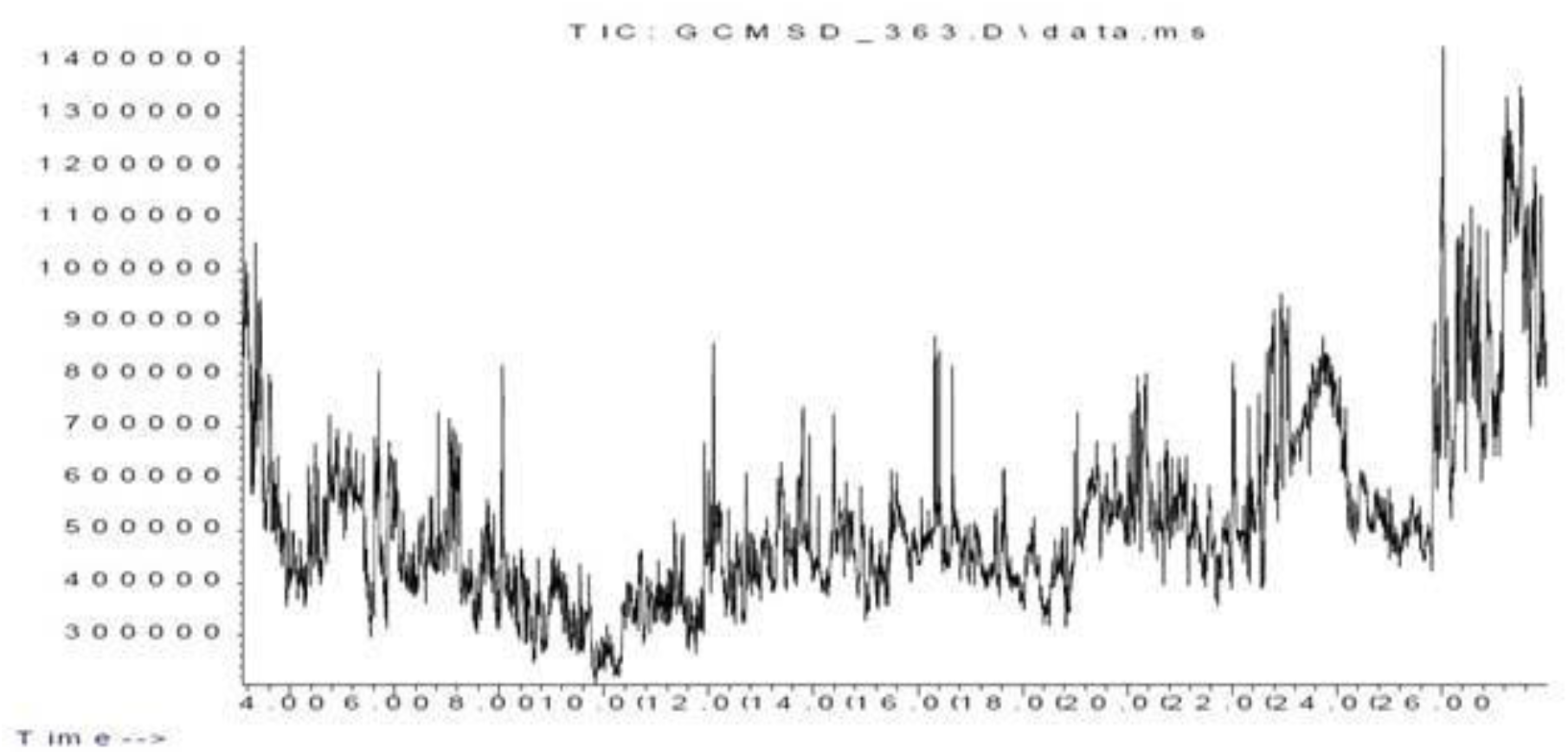
Chromatogram of *C. ambrosoides* ethanol leaf extract

*C. ambrosoides* has its highest percentage ACE inhibitory activity at 2.5mg/ml (95.990± 0.299). Also dose dependence is seen in the ACE inhibitory activity of *C. ambrosoides* with an IC50 value of 1.105mg/ml.

### 3.3. GC-MS analysis of C. ambrosoides ethanol leaf extract

The compounds identified in the extract are depicted in Table 1. A total of 92 compounds were present in the extract. The GCMS chromatogram is depicted in Figure 3.

**Table 1.**
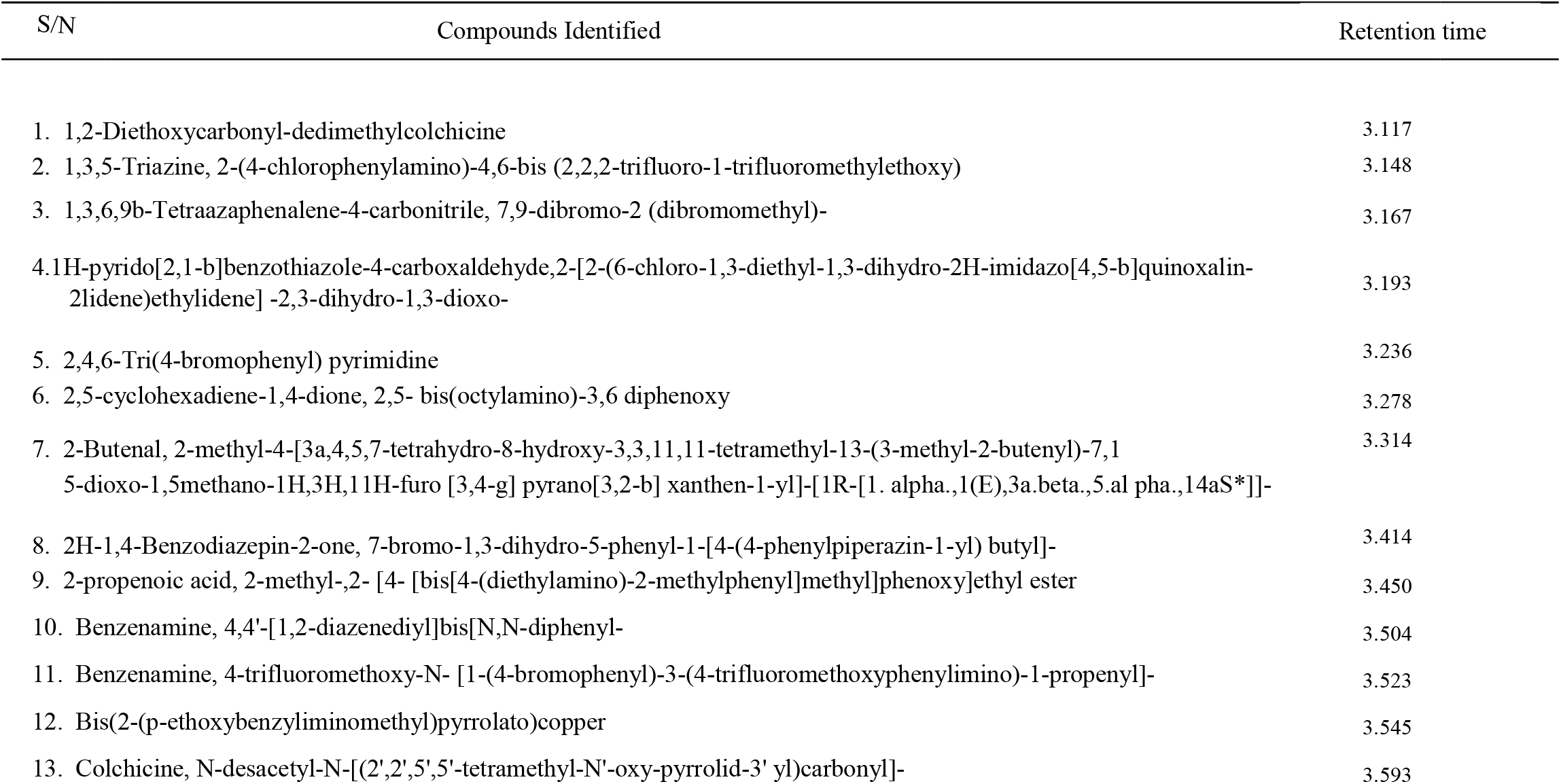
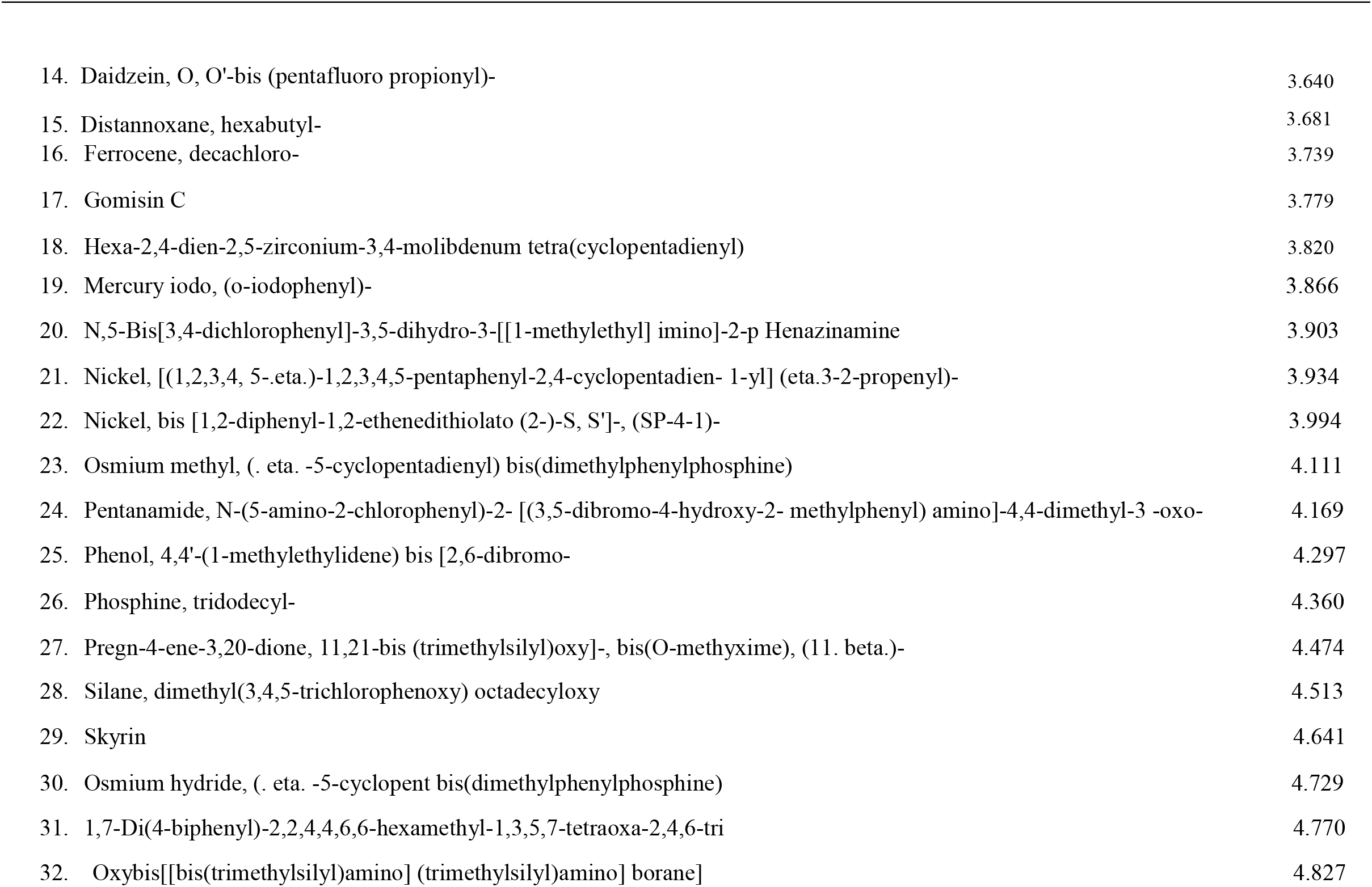
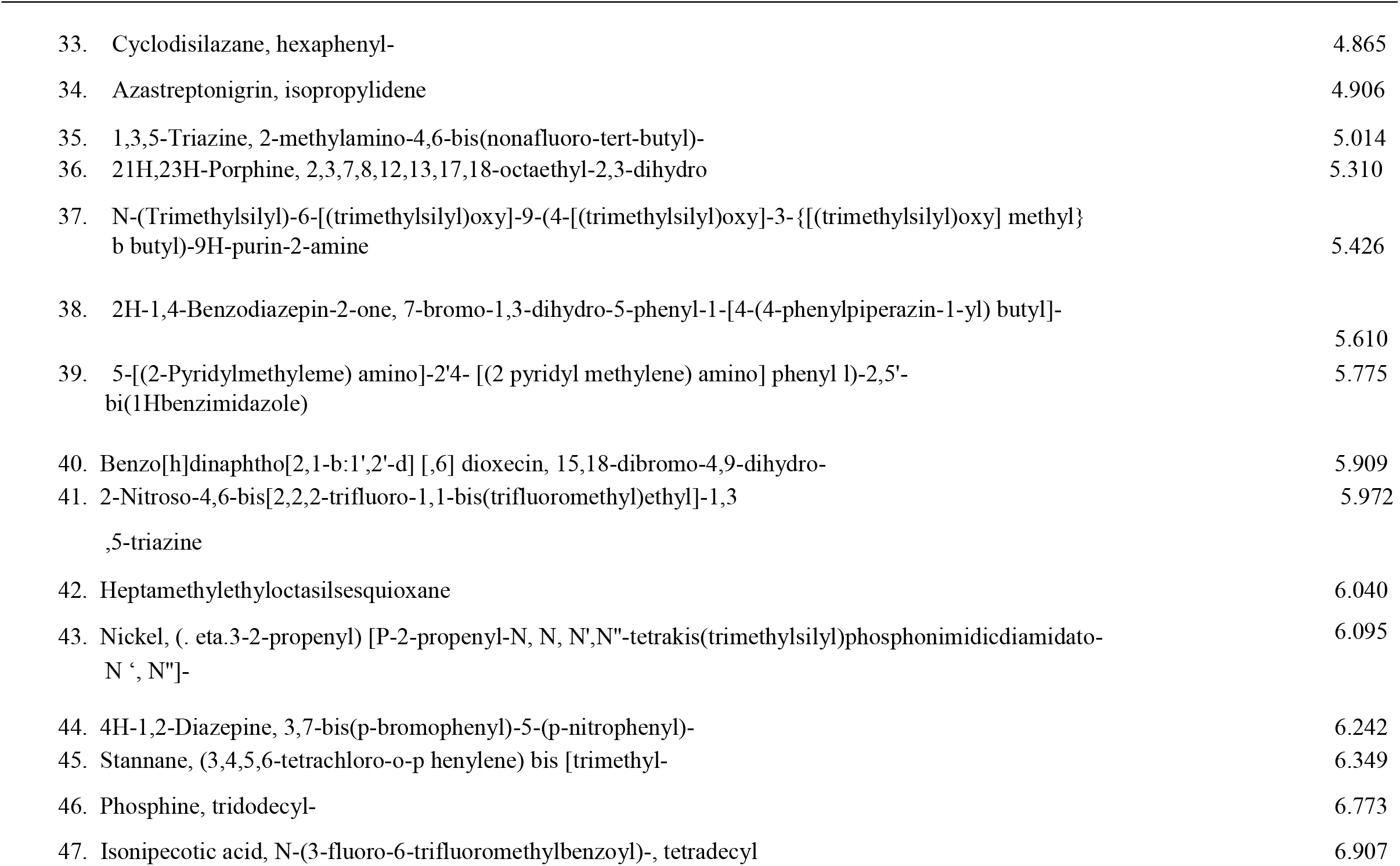
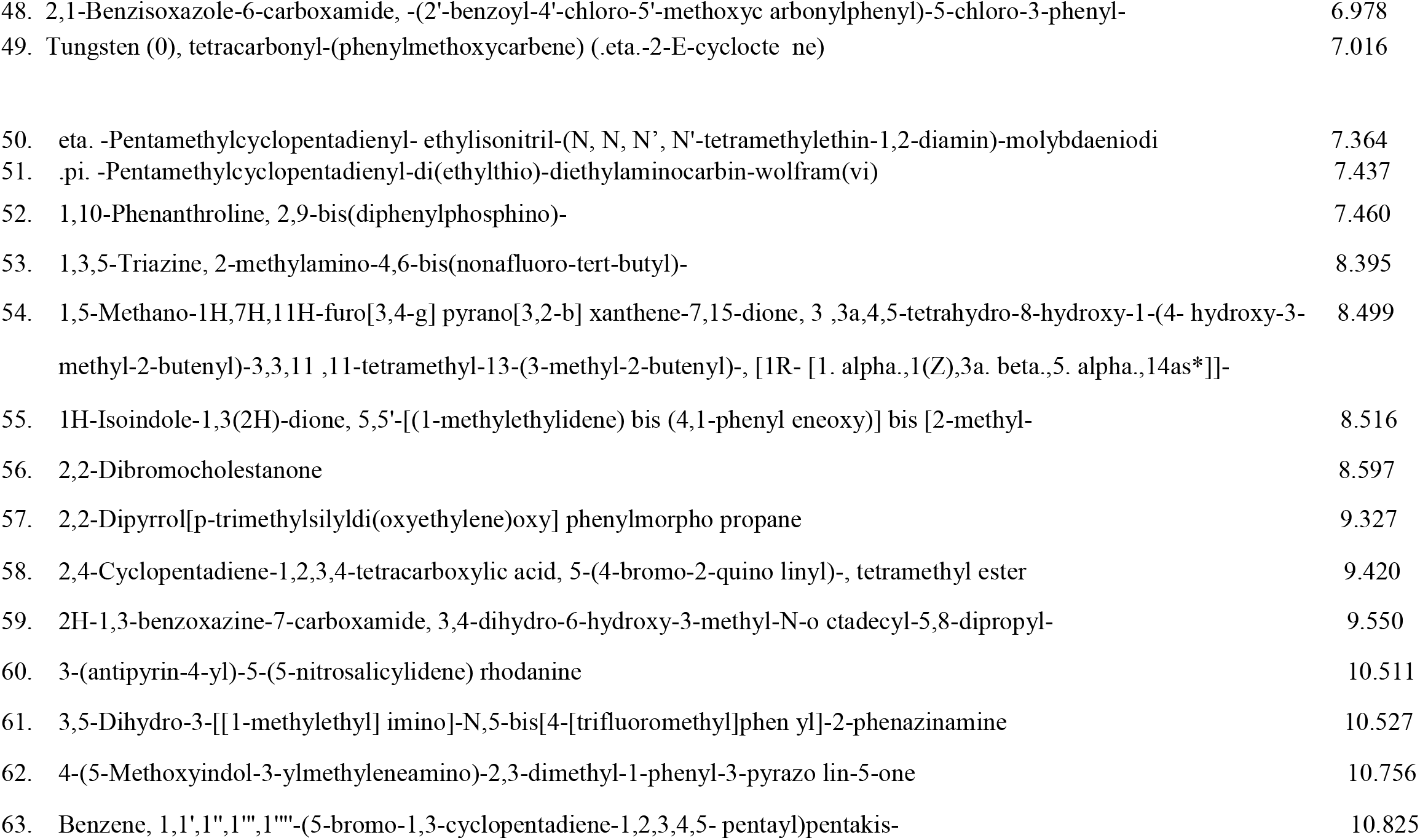
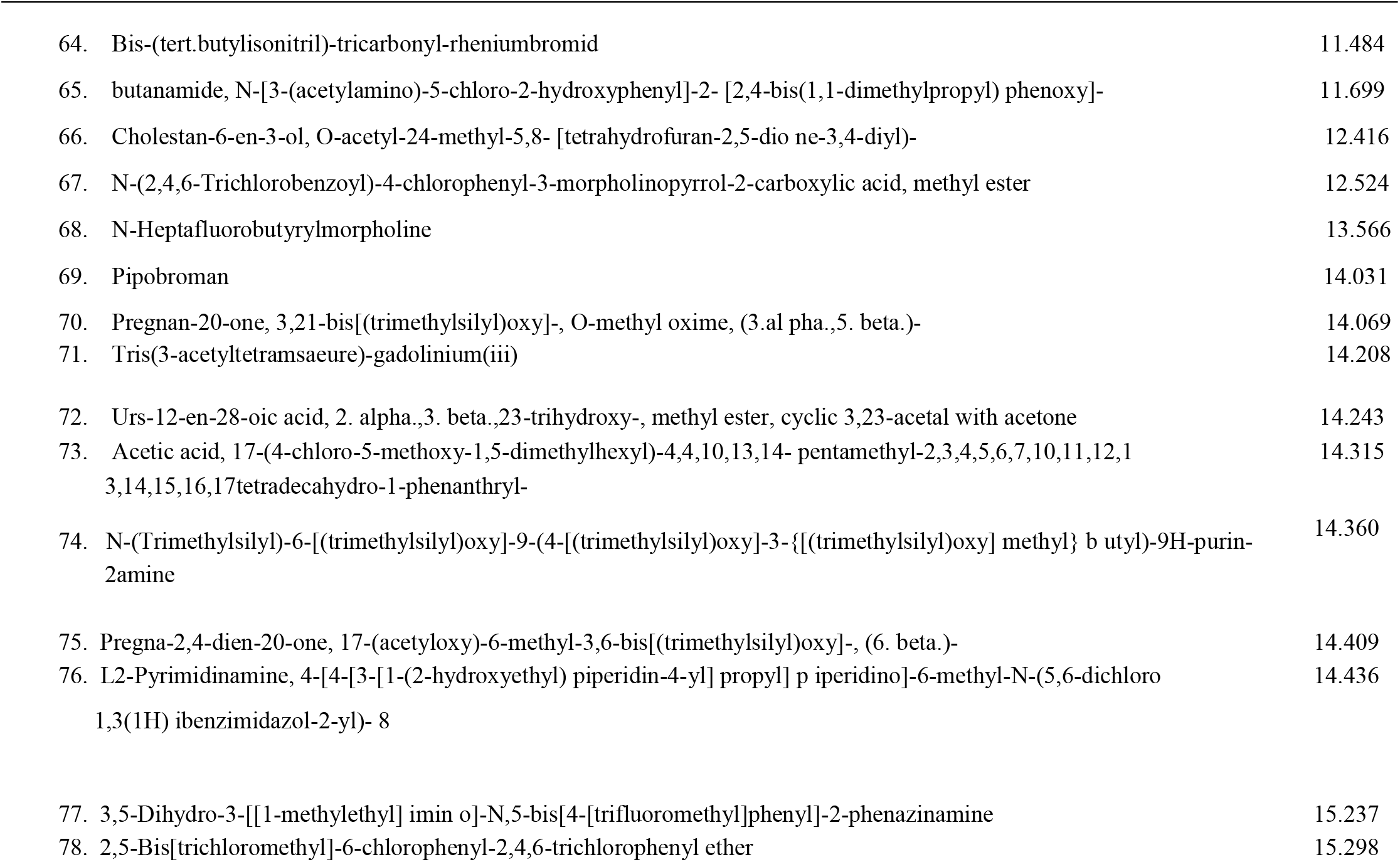
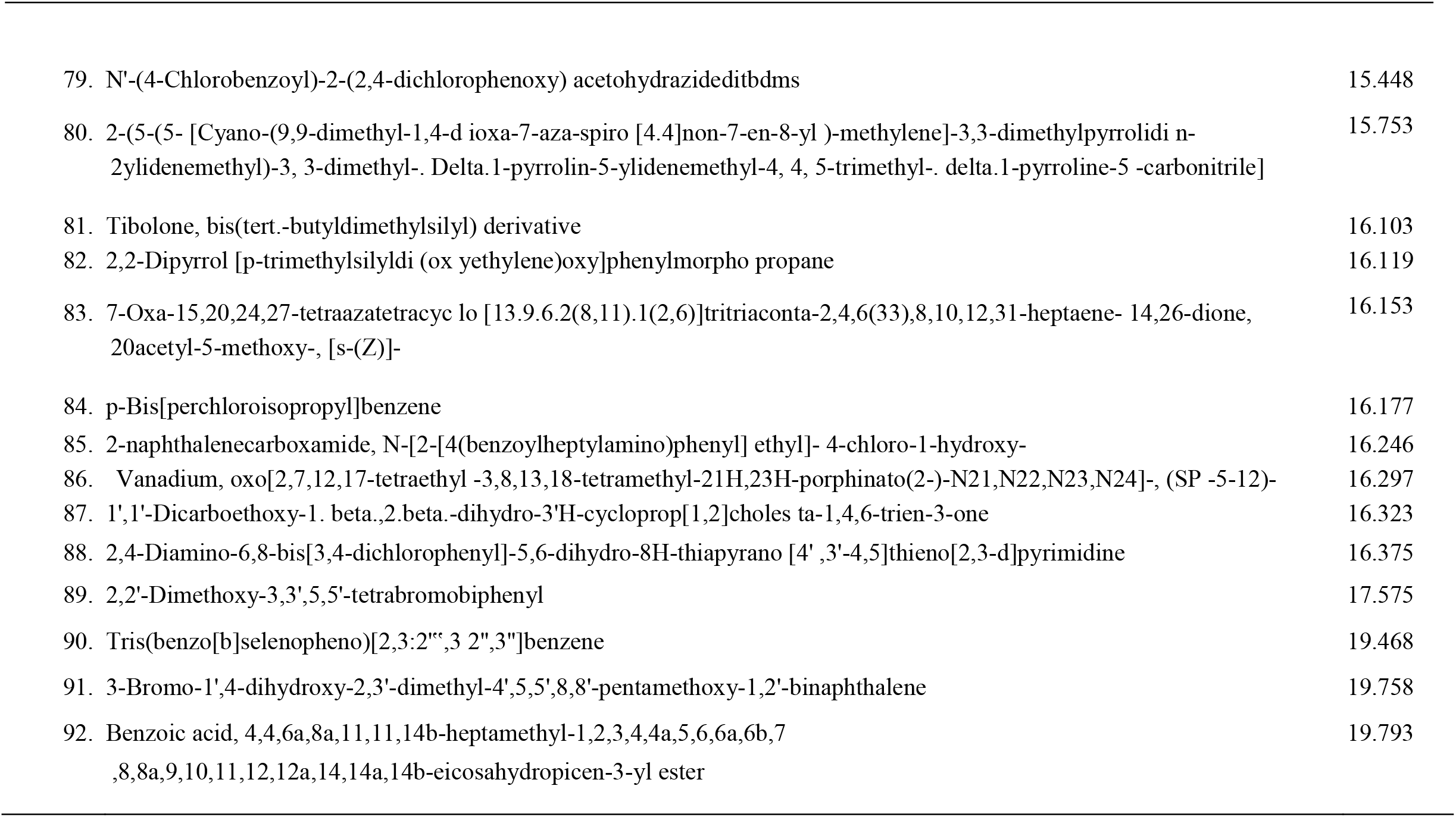
GCMS of identified compounds of C. ambrosoides ethanol leaf extract

### 3.4. In-silico studies

In-*silico* studies of the ligand-protein interaction is tabularized in Table 3.2 below showing the ligand, corresponding binding activity value, interacting amino acids and the bond types involved. From the representation, only sixteen (16) out of the ninety-one (91) phytochemicals discovered through GCMS showed computational site docking with binding affinities higher than lisinopril standard when the ligands were docked against the published active site of angiotensin converting enzyme II (ACEII). From the result, ligand 5 2,4-Diamino-6,8-bis[3,4-dichlorophenyl]-5,6-dihydro-8H-thiapyrano[4’,3’4,5]thieno[2,3d]pyrimidine showed the highest binding affinity (−8.0Kcal/mol), showing value with better site binding affinity than Lisinopril standard (6.8 Kcal/mol). Of the represented interacting amino acids in the ACEII protein binding with *C. ambrosoides* phytochemicals (ligands), GLU123, ARG124, LYS118, TRP59 and ILE88 are found to be often repetitive in interactions as seen in Figure 8.

**Table 2.**
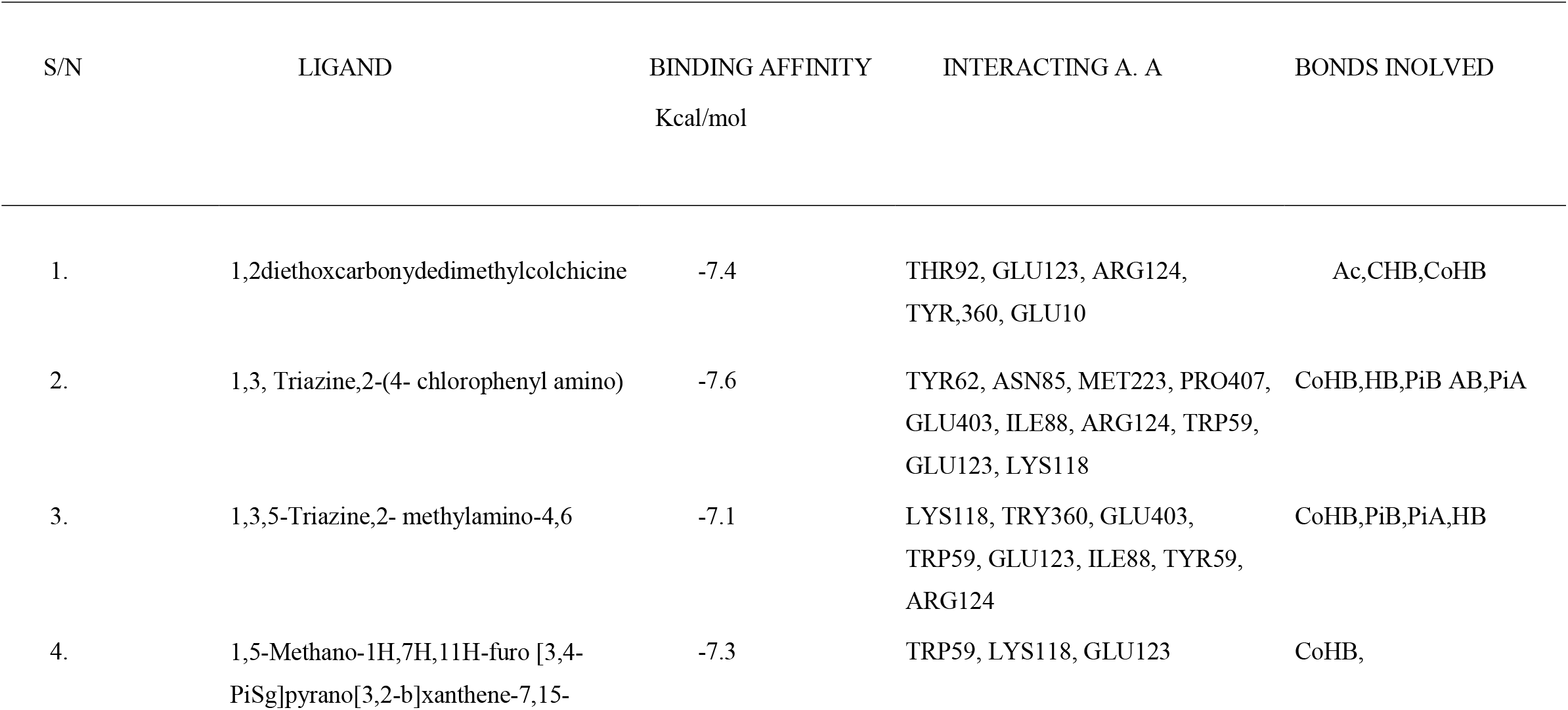
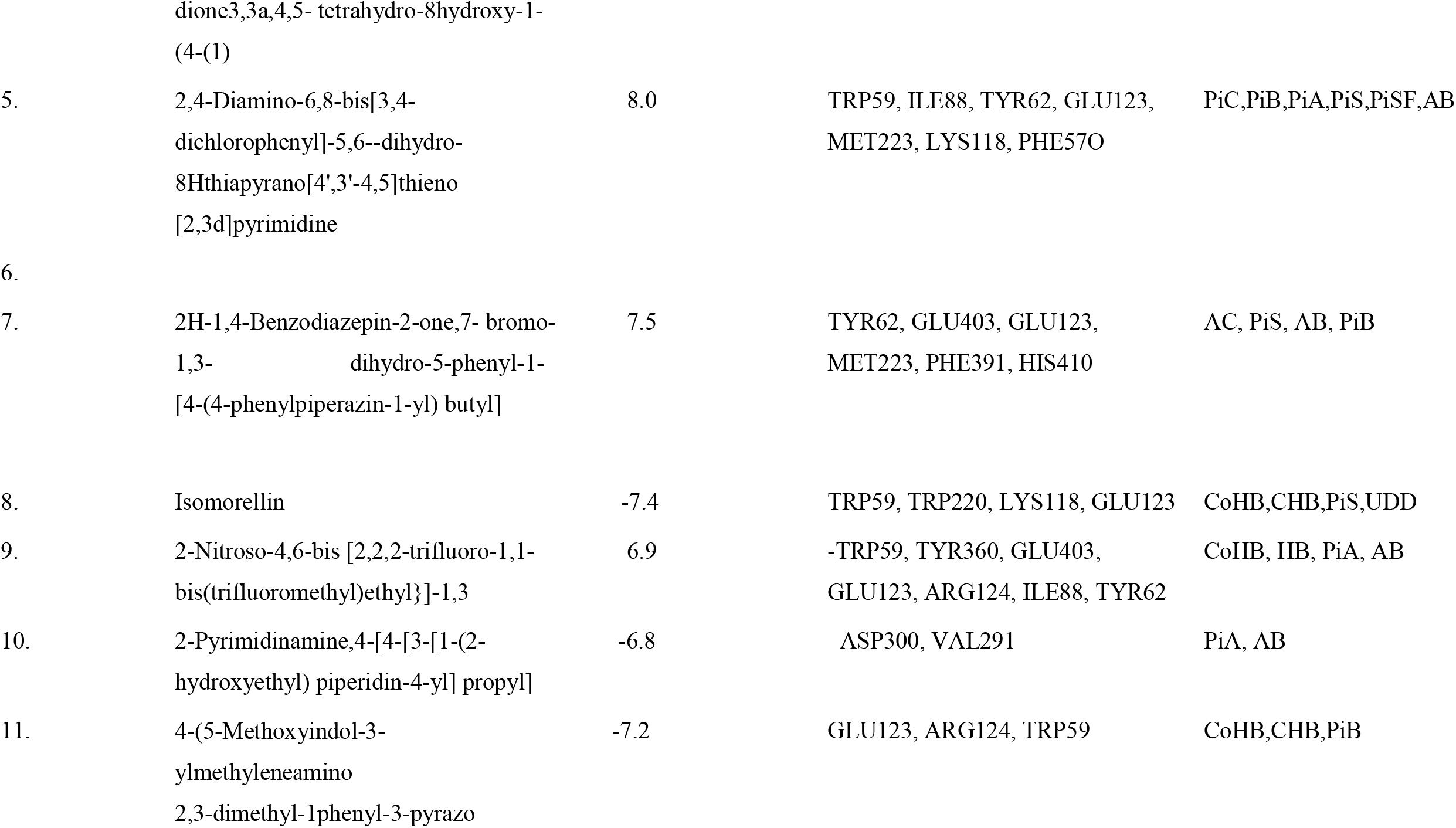
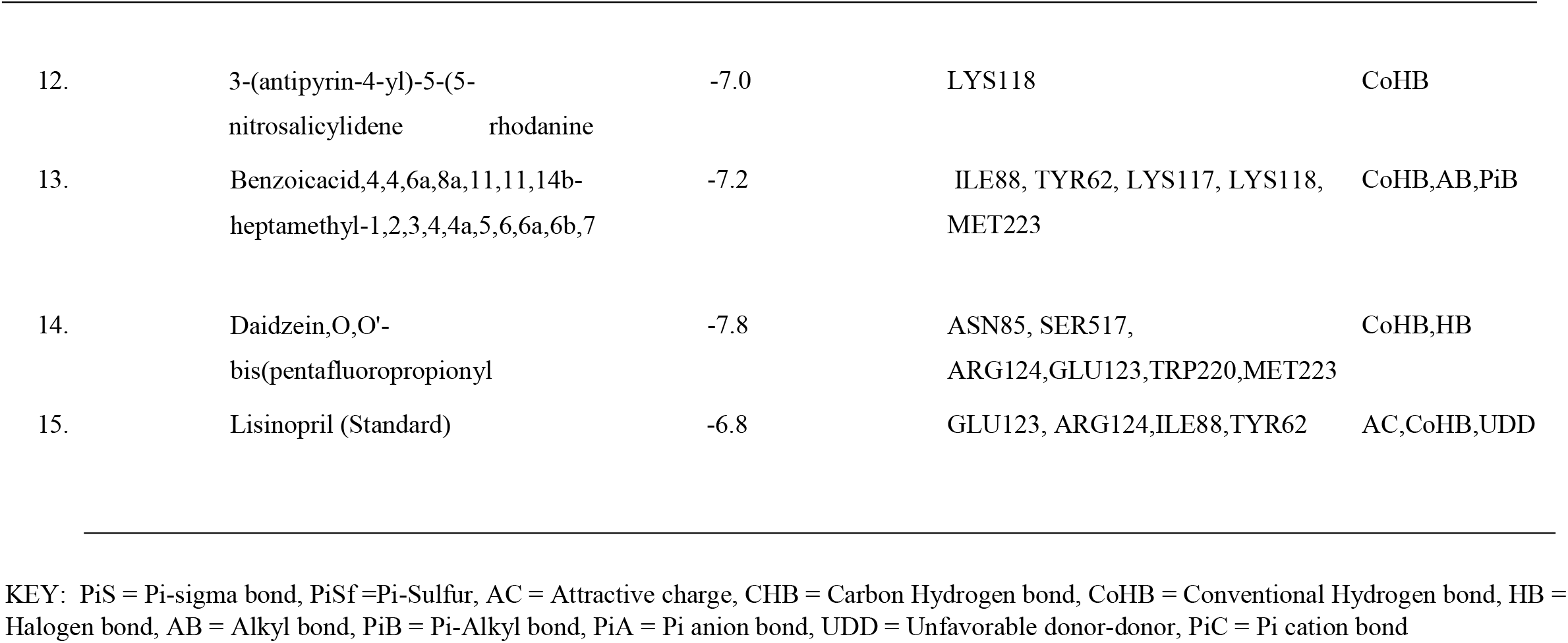
Molecular interaction of ligands with a higher binding affinity with ACE protein compared to the standard drug

| S/N | LIGAND | BINDING AFFINITY<br>Kcal/mol | INTERACTING A. A | BONDS INVOLVED |
| --- | --- | --- | --- | --- |
| 1. | 1,2diethoxycarbonyldedimethylcolchicine | -7.4 | THR92, GLU123, ARG124,<br>TYR,360, GLU10 | Ac,CHB,CoHB |
| 2. | 1,3, Triazine,2-(4- chlorophenyl amino) | -7.6 | TYR62, ASN85, MET223, PRO407,<br>GLU403, ILE88, ARG124, TRP59,<br>GLU123, LYS118 | CoHB,HB,PiB AB,PiA |
| 3. | 1,3,5-Triazine,2- methylamino-4,6 | -7.1 | LYS118, TRY360, GLU403,<br>TRP59, GLU123, ILE88, TYR59,<br>ARG124 | CoHB,PiB,PiA,HB |
| 4. | 1,5-Methano-1H,7H,11H-furo [3,4-<br>PiSg]pyrano[3,2-b]xanthene-7,15- | -7.3 | TRP59, LYS118, GLU123 | CoHB, |
|  | dione3,3a,4,5- tetrahydro-8hydroxy-1-(4-(1) |  |  |  |
| 5. | 2,4-Diamino-6,8-bis[3,4-dichlorophenyl]-5,6--dihydro-8Hthiapyrano[4',3'-4,5]thieno[2,3d]pyrimidine | 8.0 | TRP59, ILE88, TYR62, GLU123, MET223, LYS118, PHE570 | PiC,PiB,PiA,PiS,PiSF,AB |
| 6. |  |  |  |  |
| 7. | 2H-1,4-Benzodiazepin-2-one,7- bromo-1,3-dihydro-5-phenyl-1-[4-(4-phenylpiperazin-1-yl) butyl] | 7.5 | TYR62, GLU403, GLU123, MET223, PHE391, HIS410 | AC, PiS, AB, PiB |
| 8. | Isomorellin | -7.4 | TRP59, TRP220, LYS118, GLU123 | CoHB,CHB,PiS,UDD |
| 9. | 2-Nitroso-4,6-bis [2,2,2-trifluoro-1,1-bis(trifluoromethyl)ethyl}]-1,3 | 6.9 | -TRP59, TYR360, GLU403, GLU123, ARG124, ILE88, TYR62 | CoHB, HB, PiA, AB |
| 10. | 2-Pyrimidinamine,4-[4-[3-[1-(2-hydroxyethyl) piperidin-4-yl] propyl] | -6.8 | ASP300, VAL291 | PiA, AB |
| 11. | 4-(5-Methoxyindol-3-ylmethyleneamino 2,3-dimethyl-1phenyl-3-pyrazo | -7.2 | GLU123, ARG124, TRP59 | CoHB,CHB,PiB |
| 12. | 3-(antipyrin-4-yl)-5-(5-nitrosalicylidene rhodanine | -7.0 | LYS118 | CoHB |
| 13. | Benzoic acid, 4,4,6a,8a,11,11,14b-heptamethyl-1,2,3,4,4a,5,6,6a,6b,7 | -7.2 | ILE88, TYR62, LYS117, LYS118, MET223 | CoHB, AB, PiB |
| 14. | Daidzein, O,O'-bis(pentafluoropropionyl | -7.8 | ASN85, SER517, ARG124, GLU123, TRP220, MET223 | CoHB, HB |
| 15. | Lisinopril (Standard) | -6.8 | GLU123, ARG124, ILE88, TYR62 | AC, CoHB, UDD |

KEY: PiS = Pi-sigma bond, PiSf = Pi-Sulfur, AC = Attractive charge, CHB = Carbon Hydrogen bond, CoHB = Conventional Hydrogen bond, HB = Halogen bond, AB = Alkyl bond, PiB = Pi-Alkyl bond, PiA = Pi anion bond, UDD = Unfavorable donor-donor, PiC = Pi cation bond

**Figure 8.**
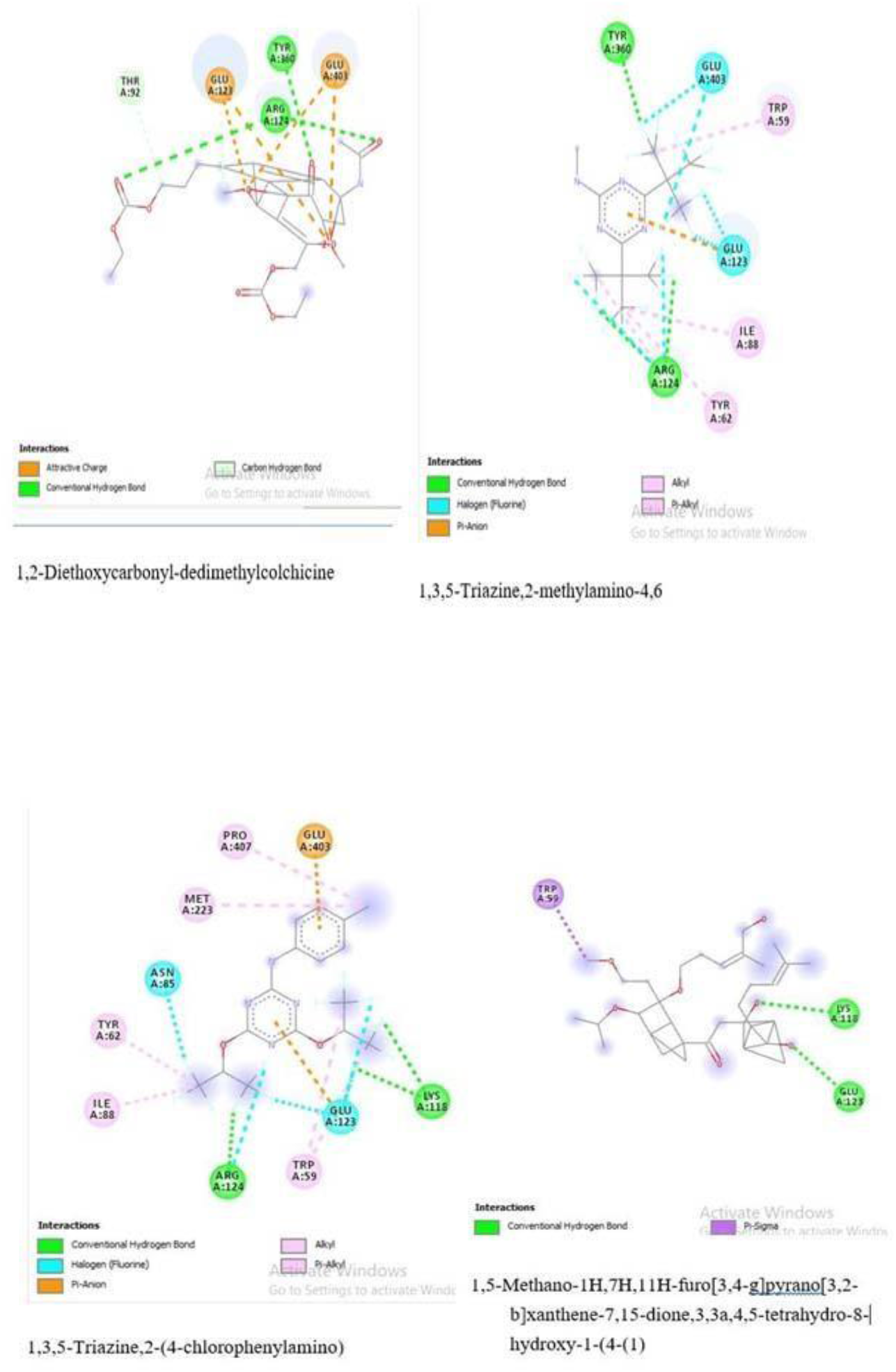
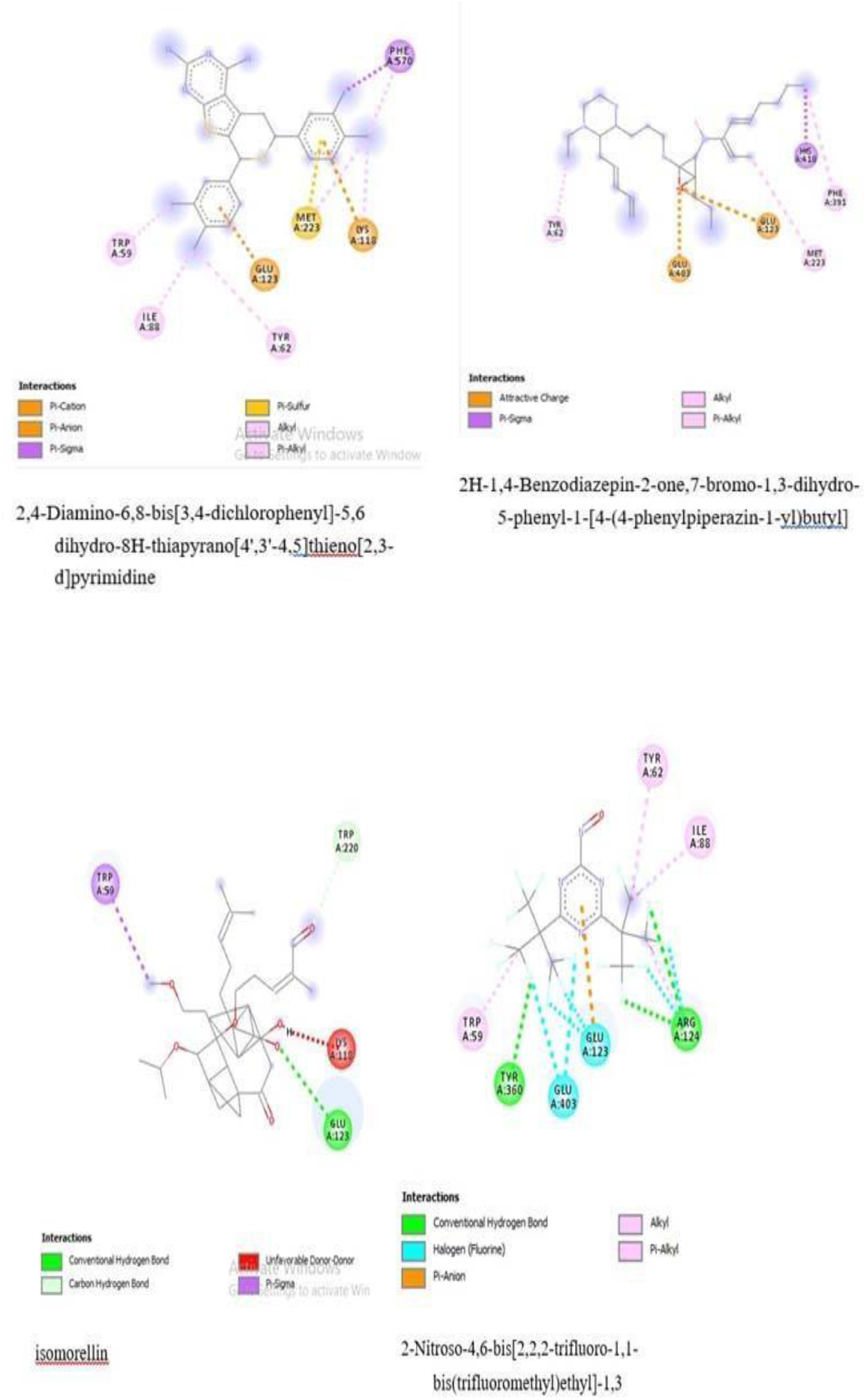
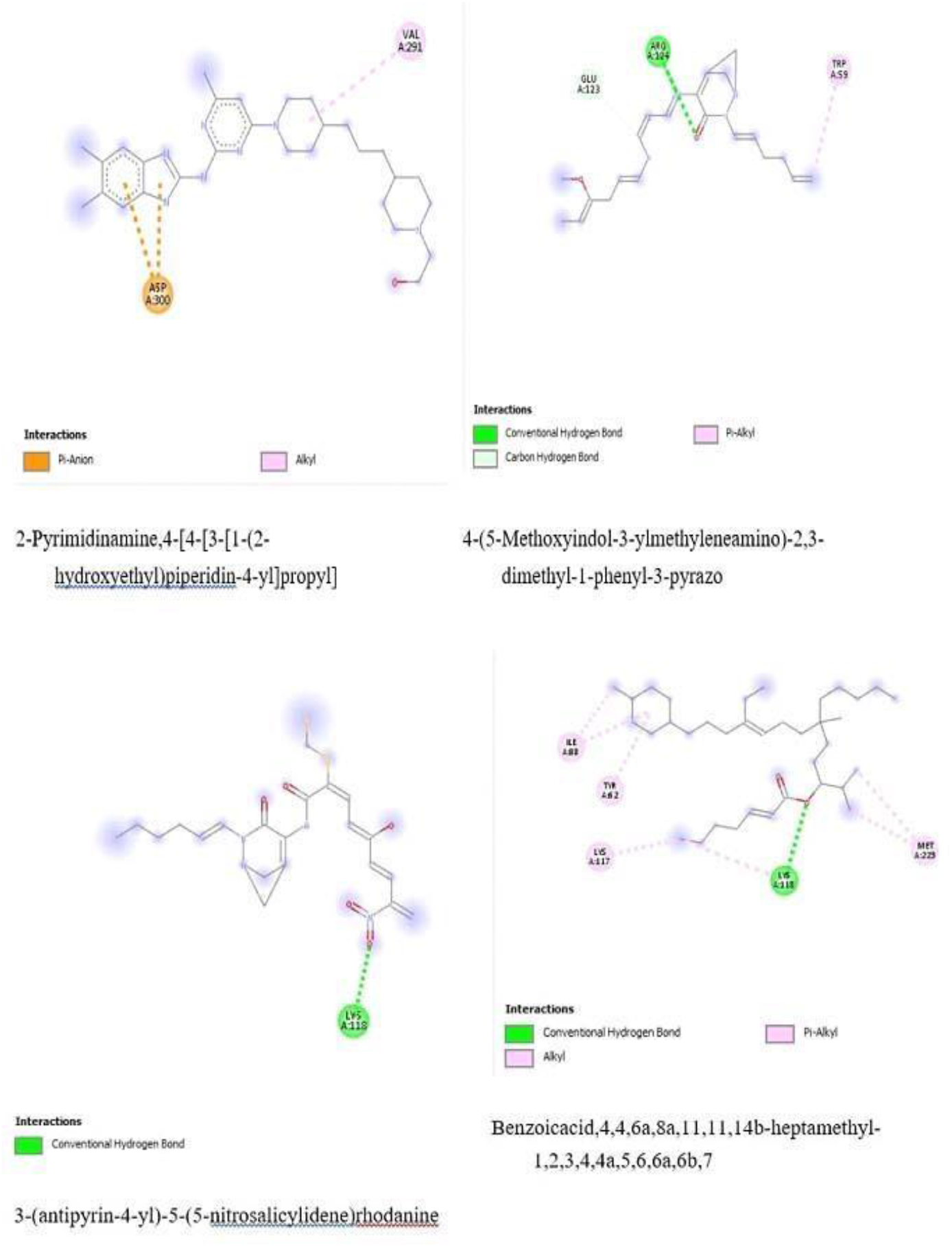
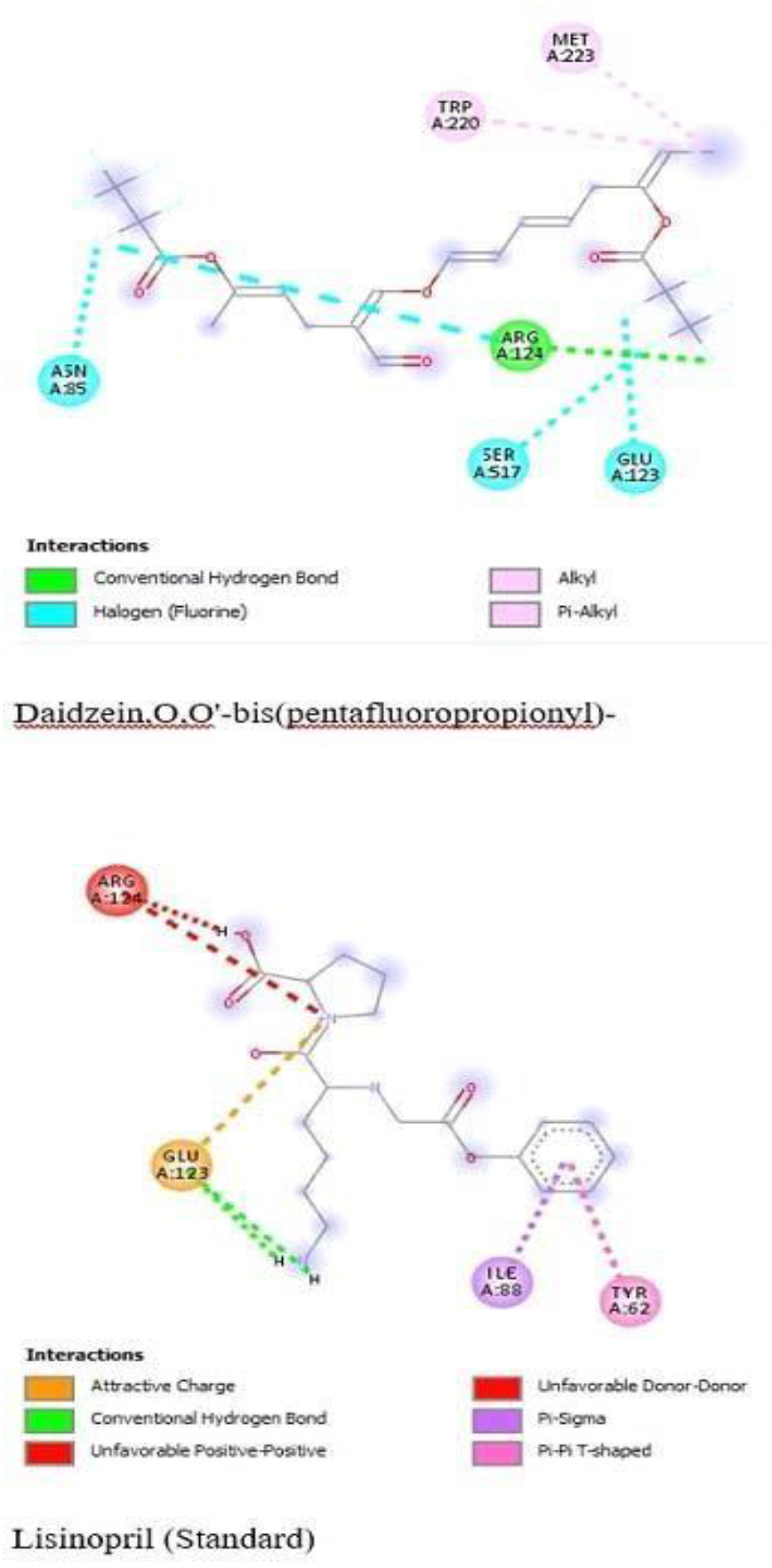
2D representation of ligand-ACE(protein) bindings.

## 4. Discussion

Oxidative stress by radicals and reactive oxygen species as superoxide anion, hydroxyl radicals, singlet oxygen etc have been indicted in the pathogenesis of various diseases ranging from malaria infection to neurological dysfunctions and degenerations to cardiovascular complications as hypertension [11]. Although intrinsic mechanisms such as antioxidant enzymes as catalase and superoxide dismutase often ameliorate physiological ROS and their developing complications, antioxidant supplements especially from plant sources cannot be overemphasized in the amelioration of oxidative stress induced ailments due to their antioxidant phytochemical constituents; chiefly polyphenols and flavonoids.

C. *ambrosoides* is an aromatic plant cultivated in different part of the world including Nigeria [12] [13], it is reported to be rich in antioxidant phytochemicals, majorly polyphenols and flavonoids [14], suggesting the discovered antioxidant potentials of the plant, thus in-*vitro* antioxidant activity of C. *ambrosoides* was evaluated via a combination of assay to verify the antioxidant potency and the mechanism. DPPH scavenging assay is a simple classical method based on the ability of extract to scavenge and quench free DPPH radicals-1,1-diphenyl-2picrylhydrazyl converting it to 1,1-diphenyl-2-picrylhydrazine or a substituted analogous hydrazine with color change through the supply of electron or hydrogen atom from the constituent antioxidants. Ethanolic extract of C. *ambrosoides* leaves induced this color change due to antioxidant bioactive compounds such as phenolic compounds and flavonoid derivatives, from which it can be inferred that the plant has the potential to quench nefarious radicals produced as superoxide anions and singlet oxygen produced during normal electron transport, accumulation of which could have result in oxidative stress. Similarly, the extract exhibited scavenging effect on nitric oxide radical (NO^•^). Nitric oxide quenching activity assay is based on the principle that sodium-nitroprusside in aqueous solution at pH of about 7.4 generates nitric oxide which act with available oxygen to form nitrite ion that can be quantified using Griss reagent, however the extract scavenges and prevent nitric oxide formation by competing with oxygen, resulting in reduced production of Nitrite [15],NO^•^ often perform a twin role as both an antioxidant and pro-oxidant depending on the relative ratios of the reactants. Antioxidant effects of NO^•^ occurs when it reacts with alkoxyl and peroxyl radical intermediates during lipid peroxidation and stabilizes the inhibition of LDL oxidation while the pro-oxidant reaction occurs when NO^•^ reacts with superoxide anion to yield peroxynitrite (ONOO^•^) a nefarious reactive nitrogen specie (RNS). The ability of Chenopodium to scavenge NO^•^ and especially its deleterious metabolite, peroxynitrite is immensely beneficial in living system as ONOO^•^ and some other NO^•^ metabolites have been indicted in various disease conditions associated with oxidative stress. Thio-barbituric acid reactive substance (TBARS) assay is an in *vitro* assay that quantifies lipid oxidation and peroxidation, based on the indirect quantification of produced malonyl-di-aldehyde which react with TBARS to form a pink chromogen. Lipid peroxidation is a nefarious consequence of accumulating free radicals and reactive oxygen species as they facilitate the oxidation of polyunsaturated fatty acids especially on plasma membrane of cells and facilitating various diseases as arthrosclerosis, autoimmune dysfunction, inflammations, cardiovascular which includes hypertension [16], one product of Lipid peroxidation is malonyl-di-aldehyde which has been implicated in protein crosslinking via lysine, histidine arginine and methionine amino acids; adduct formation with proteins and nucleic acid [16]. The Chenopodium ethanolic leaf extract inhibited TBARS reaction and chromogen formation, indicating that the extract contains antioxidant phytochemicals that can terminate lipid peroxidation and chain reaction. Ferric reducing power assay classically quantifies the ability of a suspected potential antioxidant to prevent the oxidation of metal and keep a reduced metal in its reduced state, In the present study, the extract was able to reduce the ferricyanide complex to the ferrous form, which was observed as the formation of Perl’s Prussian blue colour, indicating that the extract can function as a good electron and hydrogen atom donor which can interact with free radicals and convert them to more stable products and terminate radical chain reaction [17]. However, oxidized metals have been implicated in Fenton’s and Haber Weiss reaction which ultimately produces hydroxyl radicals which facilitates consequential nefarious reactions in the cell [17].

The result depicted from an overhaul of in-*vitro* antioxidant assays conducted all shows dose dependence in antioxidant activity, it also depicts that C. *ambrosoides* ethanol leave extract showed reduced MDA inhibitory activity when compared to ascorbate standard; the ferric reducing power exhibited by the extract healthily shows higher activity than the ascorbate standard at all concentration; the DPPH radical scavenging power shows higher activity than ascorbate standard at high concentration (2.5mg/ml and 2.0mg/ml) while at low concentrations it underperforms the standard; Nitric oxide (NO) formation inhibition assay by the extract underperforms the standard at all concentrations. A circumspect observation from the in-*vitro* antioxidant assay shows that the mechanism by which C. *ambrosoides* performs its antioxidant activity is most probably through reducing metal (e.g Fe ^3+)^ ion, and by scavenging free radicals, while other mechanisms can function as subsidiary. However, accumulation of some metal ion (Fe ^2+^, Cu^+^) has been heavily indicted in the pathogenesis of hypertension incurred from oxidative stress via Fenton and Haber mechanism [18].

GCMS result from this present research identified ninety-four (94) phytochemicals from the ethanol extract of C. *ambrosoides* (Table 1). This infers that C. ambrosoide is a repertoire of naturally occurring natural compounds, however the full pharmacological potentials of all the phytochemicals it contained may not be fully mined yet. Hence it is suggested that more ethnobotanical reports on the use of C. *ambrosoides* be succinctly followed up by classical scientific research as there may be more to uncover, C. *ambrosoides* may be a bank of an unknown long sought drug. Also GCMS result obtained indicated an already made establishment that the C. *ambrosoides* found in Nigeria often belong to the α-terpinene rich chemotype [19] as the GCMS reported terpinene derivatives from the leaves ethanol extract. In the same vein, the phytochemical compounds found in the plant has been earlier evaluated by some researchers and found a plethora of compounds, a good number of which are pharmacoactive in various respects for example ascaridole has been published to exhibit antileishmania, antiplasmodial, antiinocieptive, insecticidal, emenagouge effects and many more [13], these proving the substantial report about C.*ambrosoides* in many ethnobotanical review together with its inclusion in a national medicinal plant archive [20].

Angiotensin converting enzyme (ACE) is a sacrosanct enzyme in the RAAS mechanism facilitating hypertension pathophysiology, converting angiotensin I to angiotensin II. ACE inhibitors have been one of the potent therapeutic intervention in hypertension [21], Lisinopril being a very potent example [21]. C. *ambrosoides* ethanol extract of *chenopodium ambrosoides* leaves demonstrated ACE inhibiting potentials which is seen to be dose dependent. This finding is corroborated by computational experiment which revealed that fourteen (14) of the GCMS discovered phytochemicals of ethanol extract of C. *ambrosoides* leaves docked favorably at ACE protein binding site just as Lisinopril standard, however with binding affinities better than even the lisinopril standard. In addition, most of these ligands bind to the same amino acids lisinopril standard binds on with stable hydrogen bonds that denote protein-ligand stability, thus revealing that phytochemicals from C. *ambrosoides* can be harnessed, given much more thorough screening, optimization and designed to be a better cheap natural antihypertensive substitute to most synthetic ACE inhibitors.

It may then conclusively be inferred that the antihypertensive potential of C. *ambrosoides* may be mediated by its cumulative amelioration of oxidative stress, scavenging of free radicals and inhibition of angiotensin converting enzyme.

## 5. Conclusions

This research established the antioxidant, ACE inhibiting and antihypertensive potential of the plant, with the conclusion that the plant probably could ameliorate hypertension by simultaneously inhibiting ACE and quenching oxidative stress.

The plant may be a safer substitute for synthetic antihypertensive drugs, majority of which come with inherent nefarious contraindications. It is highly envisaged that use of *C*.

*ambrosoides* as an anti-oxidative antihypertensive drug will be more cost effective, easily accessible and provide medical solution specially to average citizens and rural dwellers who due to finance or availability could not have access to needed classical antihypertensive drug

## Acknowledgments

This research has no acknowledgement

## Conflict of Interest

The authors declare that there is no conflict of interest regarding the publication of this paper.

## Author Contributions

**Ayinde Olaniyi:** conceptualization, methodology, validation, supervision, writing-review and editing, visualization, resources.

**Oguntoye Oluwatobi**: data curation, analysis and interpretation.

**Alabi Oluwabunmi**: in-silico study

## Notes

### Competing Interest Statement

The authors have declared no competing interest.

